# SynthMLM: A framework for interpretable analysis and synthetic localisation data generation for SMLM

**DOI:** 10.64898/2026.08.12.743882

**Authors:** Louis Gall, Sandeep Shirgill, Helen Abbott, Daniel J. Nieves, Dylan M. Owen

**Affiliations:** Centre for Systems Modelling and Quantitative Biomedicine, University of Birmingham, Birmingham, B15 2TT, United Kingdom; Department of Immunology and Immunotherapy, University of Birmingham, Birmingham, B15 2TT, United Kingdom; School of Mathematics, College of Engineering and Physical Sciences, University of Birmingham, Birmingham, B15 2TT, United Kingdom; Research Software Group, Advanced Research Computing, University of Birmingham, Birmingham, B15 2TT, United Kingdom; Centre of Membrane Proteins and Receptors (COMPARE), University of Birmingham, Birmingham, B15 2TT, United Kingdom

**Keywords:** Single-molecule localisation microscopy (SMLM), Synthetic data generation, Interpretable machine learning, Bioimage analysis

## Abstract

Quantitative analysis of single-molecule localisation microscopy (SMLM) data remains challenging because biologically diverse, well-annotated datasets are limited, whilst nanoscale protein organisation is heterogeneous and difficult to describe with hand-tuned metrics. We present SynthMLM, a framework that infers interpretable structural descriptors from experimental SMLM data and uses these descriptors to generate synthetic localisation datasets. We demonstrate SynthMLM by generating descriptor-matched synthetic datasets corresponding to diverse experimental SMLM datasets and evaluating their agreement with real data using descriptor-level and embedding-based measures. By enabling controlled generation of synthetic localisation data, SynthMLM provides a practical resource for benchmarking SMLM analysis methods, testing algorithm failure modes, and developing machine-learning workflows where large, labelled datasets are required.

## 1 Introduction

The nanoscale organisation of proteins is fundamental to cellular processes, including signal transduction, membrane organisation, cytoskeletal architecture and mechanical regulation [1–3]. Single-molecule localisation microscopy (SMLM) [4] can resolve these spatial organisations at length scales inaccessible to conventional fluorescence microscopy. However, converting localisation coordinates into quantitative, biologically meaningful and generalisable descriptions remains challenging. A common limitation when applying machine learning approaches to biological imaging is the mismatch between model complexity and data availability. Although SMLM experiments can produce large numbers of localisation events, the number of biologically diverse, well-annotated datasets remains small relative to the requirements of modern machine learning. SMLM therefore occupies a difficult regime: datasets are too complex for manual analysis, but often too limited to robustly train data-intensive models. Existing approaches either prioritise interpretability, through specialised analysis pipelines [5–7] or flexibility, using deep learning that can be harder to interpret [8].

Here, we introduce SynthMLM (Figure 1), a framework that links automated SMLM analysis with synthetic data generation through interpretable structural descriptors. SynthMLM consists of two integrated components: a synthetic data generator that produces localisation patterns with user-defined fibre-like, ring-like or clustered structures, and a convolutional neural network (CNN) that estimates the corresponding descriptor distributions from experimental SMLM images. These inferred distributions can be used to generate synthetic companion datasets that reproduce the selected structural features of the original data. SynthMLM is integrated into the nano-org ecosystem [9], allowing experimental datasets to be parameterised and used as templates for synthetic data generation within an existing SMLM data resource. The model and data generation software are designed to model clustered, fibre-like [10, 11] and ring-like [12] localisation patterns, which we refer to generically as structural elements.

**Fig. 1.**
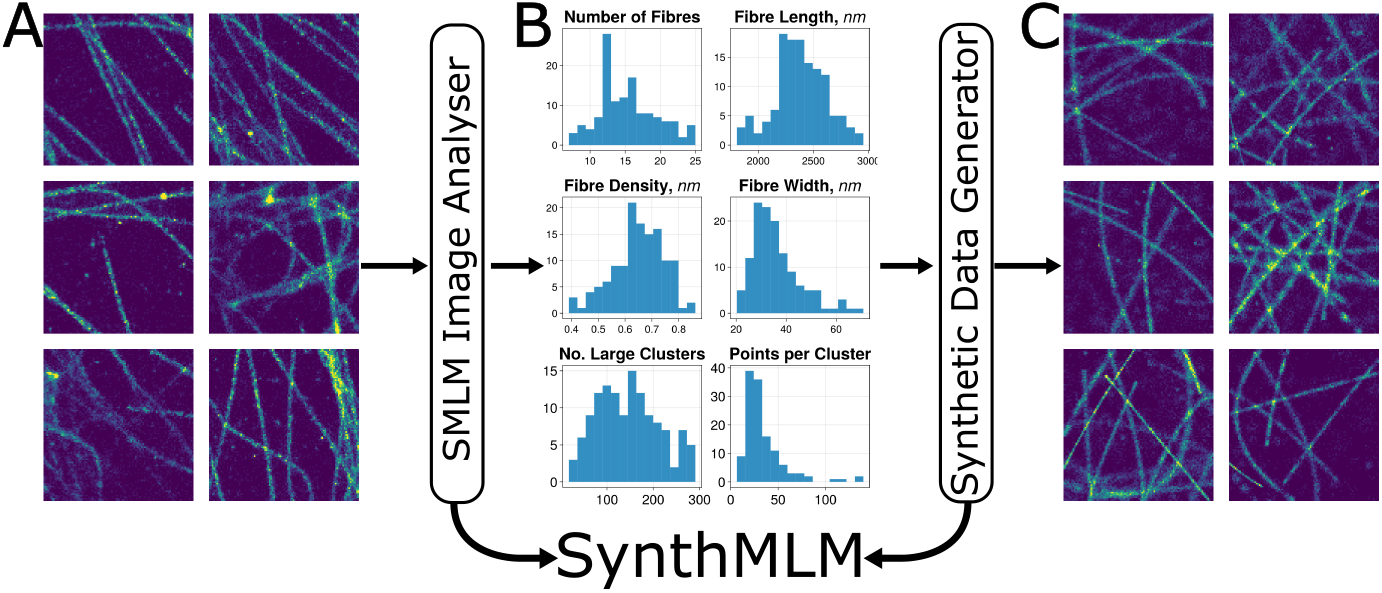
Overview of the SynthMLM workflow. A CNN-based image analyser takes rendered SMLM localisation data as input and estimates distributions of interpretable structural descriptors. These descriptor distributions can then be used by the synthetic data generator to produce new localisation patterns with matched structural properties. A) Experimental SMLM data depicting microtubules in COS-7 cells. B) Histograms showing the distributions of a selection of fibre and cluster structural descriptors inferred by the SMLM image analysis model. C) Synthetic SMLM images generated from the structural descriptor distributions inferred from the dataset in A.

This descriptor-based representation also allows controlled interpolation between measured experimental conditions. For example, inferred descriptor distributions from multiple drug doses can be interpolated to generate synthetic localisation data representing intermediate conditions.

## 2 Results

The synthetic data generator constructs SMLM-like localisation patterns by distributing localisation events around geometrically defined scaffolds that are randomly placed within the image window. Localisation placement is governed by parameterised probability distributions corresponding to interpretable structural features such as fibre length, curvature and density, ring radius and cluster occupancy. The generator was designed to cover common SMLM morphologies, including clustered, fibre-like, ring-like and mixed localisation patterns. The CNN can also be used directly as an analysis tool, estimating descriptor distributions that can be compared across experimental conditions. These descriptor distributions can then be passed to the generator to produce matched synthetic companion datasets or to interpolate between measured conditions. Each synthetic image is defined by the number of structural elements present, including fibres, rings, clusters and low-occupancy puncta, together with the descriptor distributions that define each element class.

We first applied SynthMLM as an analysis tool for experimental SMLM data. Figure 2 shows representative SMLM images of microtubules in COS-7 cells under control conditions or following treatment with 0.1 µg mL^*−*1^ or 1 µg mL^*−*1^ nocoda-zole. Figure 2B shows a subset of the output of this analysis, restricted to selected features that describe fibres and the number of larger clusters present. Consistent with nocodazole-induced microtubule disruption, increasing drug concentration was associated with fewer fibre-like structures, reduced fibre length and density, and an increased number of cluster-like structures.

**Fig. 2.**
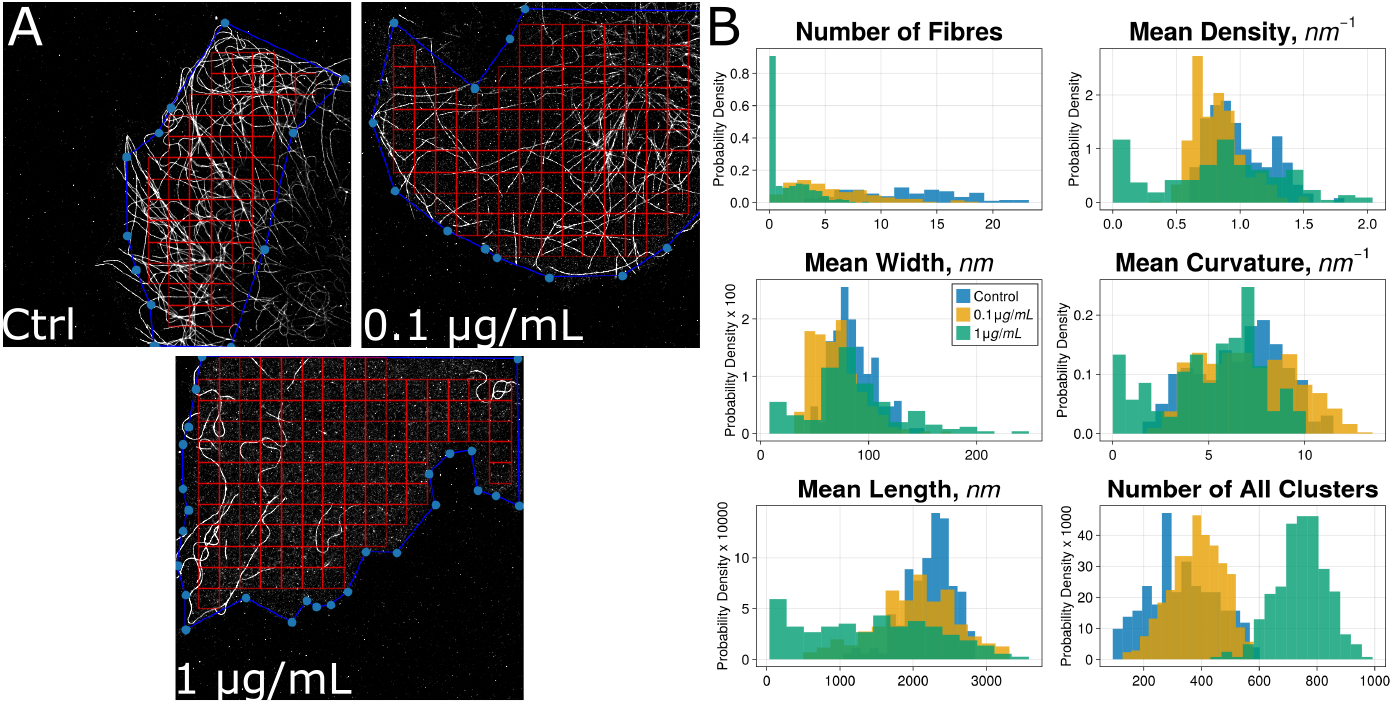
Using SynthMLM to analyse and compare the effect of three doses (control, 0.1 µg mL^*−*1^ and 1 µg mL^*−*1^) of nocodazole on microtubules. A) Whole cell image depicting microtubules in control and after treatment with nocodazole in COS-7 cells. Each image is divided into a grid of 3µm× 3µm sub-images (red lines) that are used as input data for the model. B) Histograms depicting various fibre statistics and number of clusters per sub-image for the three conditions, normalised as a probability density function to allow easy comparison between conditions.

We next used inferred descriptor distributions to generate synthetic companion datasets. For input into the deep learning model, the images were divided into 3µm× 3µm sub-images, with nine representative sub-images from each condition shown in Figures 3A-C. The inferred descriptor distributions were then used as inputs to the synthetic data generator to create a matched synthetic dataset for each condition, as depicted in Figures 3D-F for the control and two doses.

**Fig. 3.**
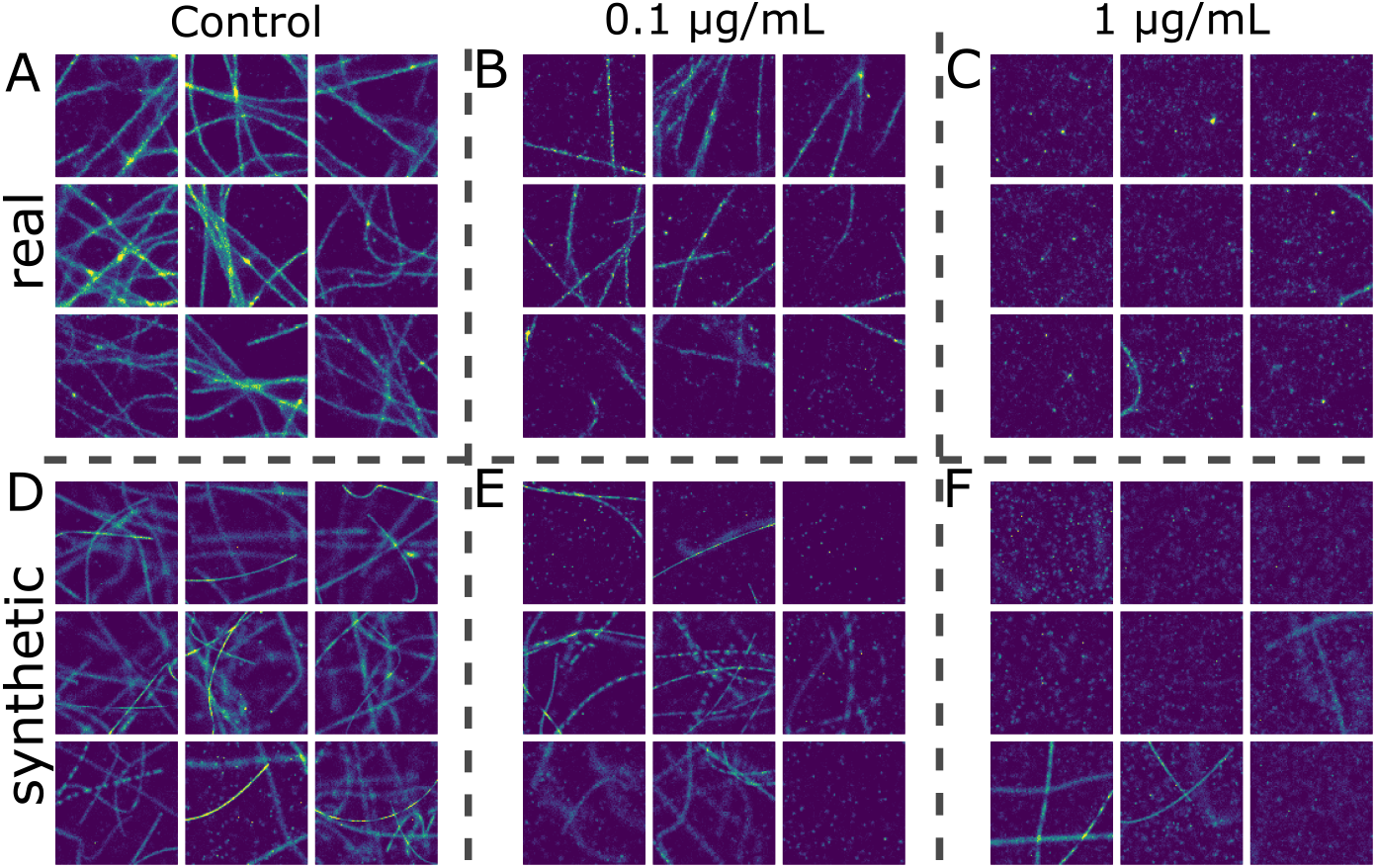
Using SynthMLM to generate descriptor-matched synthetic datasets of three different doses (control, 0.1 *µ*g mL^*−*1^ and 1 *µ*g mL^*−*1^) of nocodazole on microtubules in COS-7 cells from Figure 2. A), B) and C) real SMLM images after being divided into 3µm× 3µm sub-images and binned into 100*×*100 pixels, for the control, 0.1 *µ*g mL^*−*1^ and 1 *µ*g mL^*−*1^ respectively, each showing nine gridded images from Figure 2A. D), E) and F) synthetic SMLM images generated from the inferred structural descriptor distributions for the control condition, 0.1 µg mL^*−*1^ and 1 *µ*g mL^*−*1^ respectively.

Visual comparison suggested that the synthetic datasets reproduced key features of the corresponding experimental data. We therefore quantified agreement between experimental and synthetic datasets using dissimilarity scores and contrastive learning embeddings [8]. Figure 4A shows the mean dissimilarity score between each nano-org dataset and its corresponding synthetic companion dataset comprising 500 synthetic sub-images. Dissimilarity was quantified using a normalised Kolmogorov-Smirnov test [13] to compare the per-pixel intensity distribution of pairs of images, as implemented in nano-org [9]. Under this metric, scores below 1 indicate that two datasets are not distinguishable at the chosen significance threshold. Figures 4B-D show inter- and intra-condition similarity scores between pairs of images normalised as probability density functions to compare datasets of different size. In each example, both the inter- and intra-condition comparisons are highly similar for the real datasets and their matched synthetic datasets. These comparisons suggest that the synthetic companion datasets reproduce key within- and between-condition relationships captured by the selected similarity metric. Figures 4E and 4F show the similarity of the contrastive learning embeddings of 3 datasets containing fibres (Figure 4E; the nocodazole exposed datasets from Figure 2 and 3) and three clustered datasets from nano-org (Figure 4F). In all cases, the synthetic datasets occupied similar regions of embedding space to the corresponding experimental data.

**Fig. 4.**
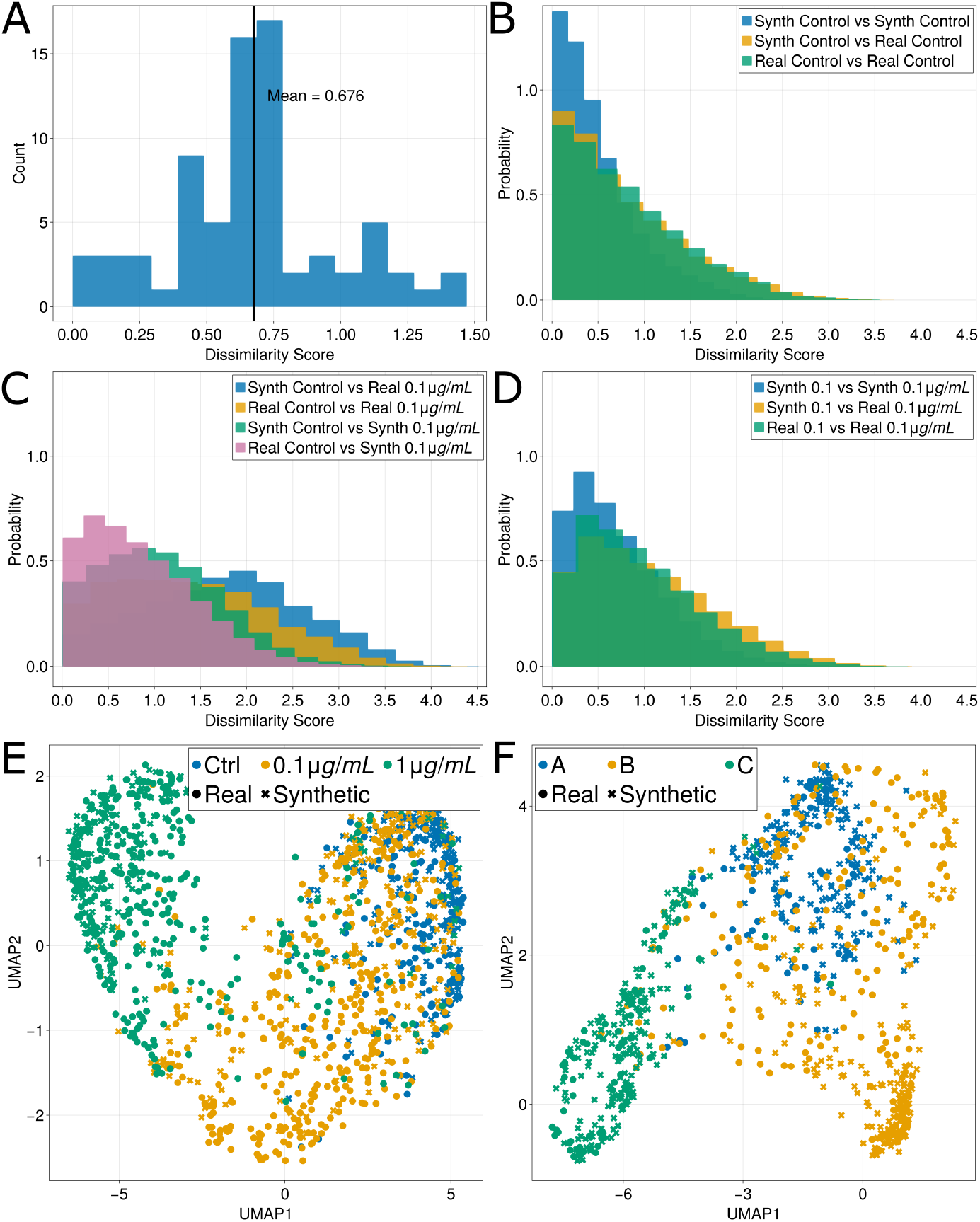
Agreement between experimental and matched synthetic datasets. A) Histogram of dissimilarity scores of every dataset on nano-org (as of 1st May 2026) and its corresponding matched synthetic dataset. A score greater than 1 implies the distributions of the real and synthetic datasets are significantly different. B) Normalised histograms showing the dissimilarity score between pairs of images from the nocodazole control from Figure 2 and 3 and its matched synthetic dataset. C) Normalised histograms showing the dissimilarity score between pairs of images from the nocodazole control and 0.1 *µ*g mL^*−*1^ nocodazole condition and their matched synthetic datasets. D) Normalised histograms showing the dissimilarity score between pairs of images from the 0.1 *µ*g mL^*−*1^ nocodazole condition and their matched synthetic datasets. E) UMAP plot of contrastive learning embedding of three datasets (circles) and their matched synthetic datasets (crosses) showing control, 0.1 *µ*g mL^*−*1^ and 1 *µ*g mL^*−*1^ nocodazole on microtubules. F) UMAP plot of contrastive learning embedding of three cluster-based datasets (circles) and their matched synthetic datasets (crosses) showing A: Lck in Jurkat E6.1 cells activated with anti-CD90 for 2 minutes, B: KIR2DL1 in NK cells and C: SLP76 in Jurkat E6.1 cells activated with anti-CD3CD28 for 5 minutes. All histograms were normalised as probability density functions to compare datasets of unequal size.

Having established agreement between real datasets and their synthetic companions using both dissimilarity scores and contrastive embedding, the third application allows for interpolation between experimental conditions, for example, estimating descriptor distributions for intermediate drug doses using existing data (Figure 5). This allows visualisation of intermediate experimental conditions and provides descriptor-level estimates through interpolation between measured doses, which are evaluated here against an experimental 0.5 µg mL^*−*1^ dataset. Using the relationships between multiple doses, we can formulate an estimated distribution of the model output parameters for an unseen dose (Figure 5A) and generate data based on these distributions (Supplementary Methods).

**Fig. 5.**
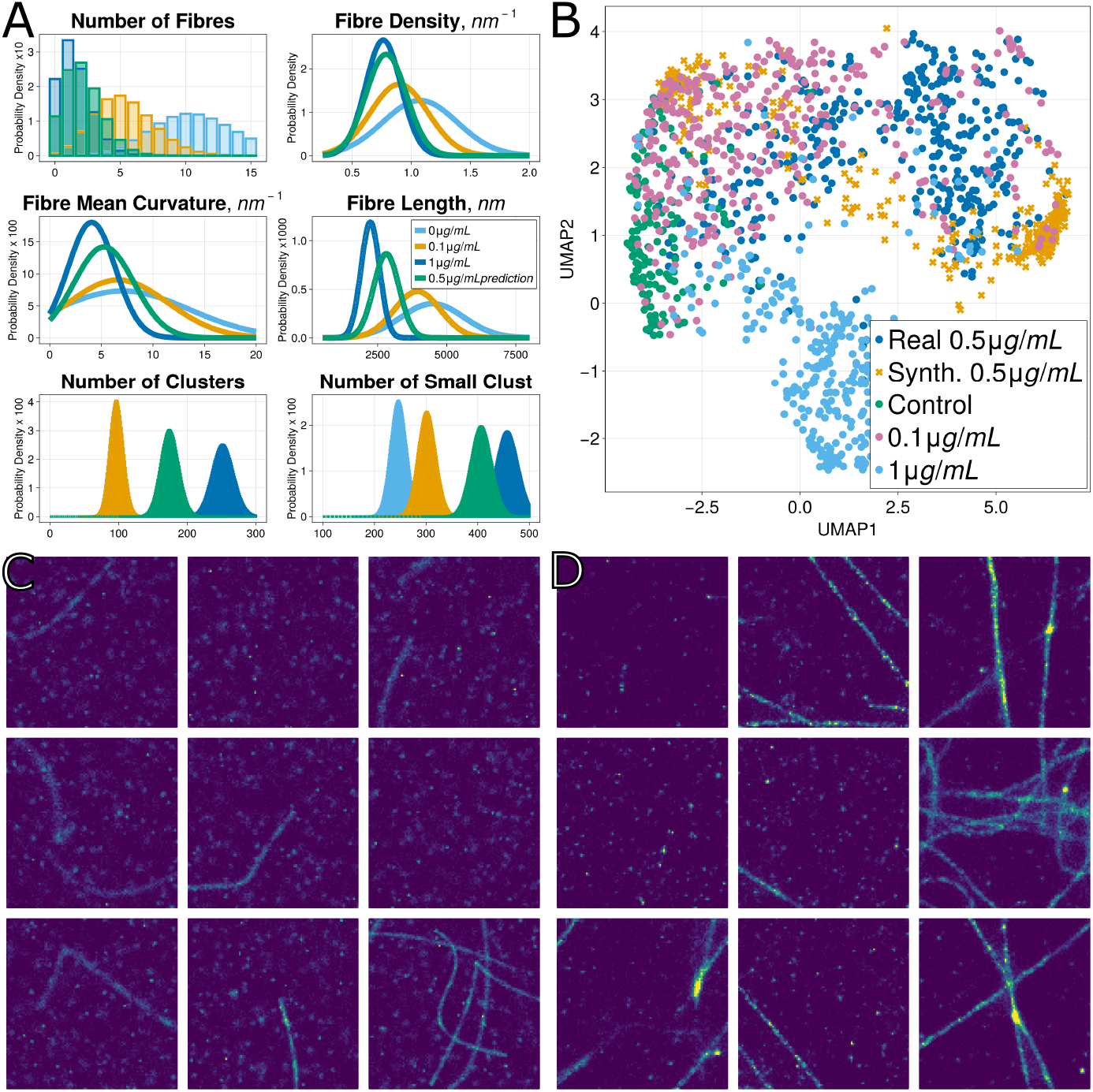
Demonstration of interpolating doses using output structural descriptors. A) Probability density functions of selected quantities from SMLM images of microtubules treated with 0 *µ*g mL^*−*1^ (light blue), 0.1 *µ*g mL^*−*1^ (yellow) and 1 *µ*g mL^*−*1^ (dark blue) of nocodazole, and interpolated values of a 0.5 *µ*g mL^*−*1^ dose of nocodazole (green). B) UMAP plot of contrastive learning embedding of an experimental 0.5 *µ*g mL^*−*1^ nocodazole dataset (blue circles), the synthetic 0.5 *µ*g mL^*−*1^ data (yellow crosses), control (green circles), 0.1 *µ*g mL^*−*1^ (pink circles) and 1 *µ*g mL^*−*1^ condition (light blue circles). C) Synthetic images of the predicted 0.5 *µ*g mL^*−*1^ exposure. D) Real SMLM images of the microtubules exposed to 0.5 *µ*g mL^*−*1^ of nocodazole in COS-7 cells.

In the contrastive learning embedding, the interpolated 0.5 *µ*g mL^−1^ synthetic dataset occupied a similar region of UMAP space (Figure 5B) to the corresponding experimental 0.5 µg mL^*−*1^ dataset. The interpolated synthetic data showed reduced variability relative to the experimental data, consistent with the smoothing imposed by interpolation between descriptor distributions. Figure 5C shows hypothetical synthetic localisation patterns for the interpolated condition that qualitatively resembled the corresponding experimental data (Figure 5D) obtained from COS-7 cells exposed to 0.5 *µ*g mL^*−*1^ of nocodazole.

## 3 Discussion

We have presented SynthMLM, a framework for analysing SMLM data and generating synthetic localisation datasets using interpretable structural descriptors. We demonstrated three principal applications of SynthMLM: analysis of real datasets, generation of synthetic companion datasets and interpolation of intermediate experimental conditions. Note that SynthMLM generates data mimicking experimental SMLM data, not distributions mimicking the underlying ground truth of the positions of proteins in the biological sample.

SynthMLM enables rapid generation of large numbers of controlled synthetic localisation datasets sampled from inferred descriptor distributions. These datasets are not a substitute for independent biological experiments. Instead, they provide labelled, parameter-controlled samples for benchmarking analysis workflows, testing failure modes and developing machine-learning methods. In contrast to fully black-box generative approaches, such as generative adversarial networks [14, 15], variational auto-encoders [16] or diffusion models [17, 18], SynthMLM generates data through explicit structural descriptors. By unifying interpretable modelling with data generation, this work provides a practical and conceptually distinct alternative to purely deep learning approaches, and provides an important tool for data-efficient machine learning in nanoscale biological imaging, grounded in interpretable structural quantities.

## 4 Methods

SynthMLM combines a synthetic data generator of SMLM localisation patterns with a convolutional neural network that analyses SMLM localisation patterns in terms of interpretable biophysical features. The synthetic data generator builds SMLM images from structural elements (clusters, rings and fibres) parameterised by user-defined distributions of the structural descriptors, while the CNN estimates the distributions of the structural descriptors governing these elements from SMLM images. Inferred distributions from experimental data can be used to generate synthetic companion datasets matching the structural properties of the original data. The Supplementary Information provides further implementation details of all aspects of SynthMLM.

The data generation software and the SMLM image analysis model were both written in the Julia programming language [19] using the Flux.jl machine learning library [20, 21]. SynthMLM is integrated into nano-org for the analysis and generation of synthetic datasets.

### Synthetic Data Generation

Synthetic SMLM images are generated by combining three structural element classes: clusters, rings and fibres. Each element is defined by a set of structural descriptors corresponding to interpretable biophysical quantities. The data generation software can sample from arbitrary probability distributions for all structural descriptors. Throughout this work, we typically used truncated normal distributions.

Clusters are represented as Gaussian point clouds with independently sampled centre locations, particle numbers and spatial widths. Two cluster populations are supported: small clusters (≤10 particles per cluster, representing diffuse low-occupancy puncta) and larger localisation clusters (> 10 particles per cluster), each with independent parameter distributions. The two classes of clusters were modelled separately to better represent experimentally observed background and heterogeneous protein organisation.

Rings are defined by a circle with a fixed radius, with particles distributed around this circle according to the density and width of each ring. Ring centres are initially placed randomly within the image window and subsequently relaxed using a physics simulation to eliminate overlaps. The ring density is defined as the linear particle density along the circumference of the circle that defines the ring. The ring width refers to the spatial extent normal to the circle that defines the ring, with a particle’s transverse position sampled from a normal distribution with standard deviation equal to half this width.

Fibres are made by placing particles around a Bézier curve [22] scaffold. The structural descriptors defining a fibre are its length, width, density, mean curvature, and two parameters describing intensity heterogeneity: the spacing between intensity peaks and the width of these intensity peaks. The density of a fibre is the linear density along the tangent direction of the Bézier curve that defines the fibre. The fibre width refers to the spatial extent normal to this curve, with a particle’s transverse position distributed according to a normal distribution with standard deviation equal to half the width of the fibre.

After all structural elements are generated, particles lying outside the image window are discarded. The synthetic data generator can output either a .csv file containing the coordinates of each simulated particle or a binned 100 × 100 image representing localisation counts within a 3µm × 3µm field of view.

### Convolutional neural network

The image analysis model is a RegNetX-based convolutional neural network [23] containing approximately 27 million parameters. The network performs multi-output regression, predicting the number of each structural element present in an image together with the mean and variance of the probability distribution associated with each structural descriptor. These outputs correspond directly to the parameters used by the synthetic data generator, providing an interpretable representation that links experimental SMLM images to synthetic data generation. In total, the network predicts the number of fibres, rings, clusters and low-occupancy puncta, and the mean and variance of 13 structural descriptors describing these elements, giving 30 output variables (Supplementary Table 1).

### Structural Descriptor Inference

The model accepts 3µm ×3µm SMLM localisation patterns binned into 100× 100 images. Input images are normalised by clamping pixel intensities to a maximum of 40, before scaling intensities to the range [0, 1]. Model outputs are transformed from the min-max normalised training space back into their corresponding physical units before further analysis.

During inference on experimental data, structural descriptor estimates are produced independently for each image. Dataset-level descriptor distributions are then obtained by aggregating these predictions across all images.

### Model Training

Using the synthetic data generator, we created a training set consisting of 10 million synthetic SMLM images containing diverse combinations of clusters, rings and fibres. To improve model generalisation, the training set encompassed a broad range of synthetic morphologies, including hypothetical or experimentally unobserved arrangements. For each generated image, structural descriptors were sampled from truncated normal distributions whose means and variances were themselves sampled over a wide range of values. The training data also included arbitrary combinations of the three structural element classes (e.g. images containing both clusters and fibres, rings and fibres, or all three element types). Such mixed morphologies are biologically relevant; for example, nocodazole treatment fragments microtubules into a mixture of shortened fibres and cluster-like structures.

Counts of each structural element and the mean and variance of the distributions of each structural descriptor were recorded for every generated image in the training dataset and used as regression targets. These targets were min-max normalised during training.

The model was trained using the AdamW optimiser [24] with a OneCycle learning-rate schedule [25] and a mean squared error loss. Training was performed for 20 epochs, with model selection based on validation performance using a held-out set of 100,000 synthetic images generated independently using SynthMLM.

### Synthetic dataset reconstruction

To generate synthetic datasets matching an experimental SMLM dataset, the trained network is applied to each image to estimate element counts and distributions of the structural descriptors.

Two reconstruction strategies were implemented. In the dataset-level approach, descriptor estimates from all images are aggregated to form global distributions for each structural descriptor. Synthetic images are then generated by sampling these distributions and passing the sampled descriptors to the synthetic data generator. This approach was used throughout, except when generating the descriptor-matched synthetic counterparts of the nano-org datasets shown in Figure 4.

The second image-level approach preserves local heterogeneity by randomly selecting individual experimental images and directly using the predicted parameter set to generate each synthetic image. This image-level reconstruction better reproduces multimodal or spatially heterogeneous datasets, where averaging structural descriptor distributions across all images would obscure biologically meaningful variation.

## Supplementary information

The online version contains supplementary material, including Supplementary Methods, Supplementary Figures, Supplementary Tables and additional references.

## Acknowledgements

The research described in this paper was carried out with the assistance of Advanced Research Computing at the University of Birmingham. This included support from the Research Software Group to convert the original academic code into a reusable Julia package, support with the development of the database and website, data storage on the Research Data Store, and computations on the BlueBEAR HPC service.

## Author Contributions

LG and DMO conceived the work. LG, SS, DJN and HA provided code and/or data and/or performed analysis. LG and DMO wrote the manuscript.

## Data Availability

All SMLM datasets analysed in this study were obtained from nano-org [9] except the Xenopus nuclear pore complex (NPC) datasets used in the Supplementary Information which were obtained from ShareLoc.XYZ via Zenodo (https://zenodo.org/records/7185997 and https://zenodo.org/records/7182227).

## Code Availability

SynthMLM is integrated into nano-org [9] for SMLM dataset analysis and structural descriptor-matched synthetic data generation. The SynthMLM software will be released as an open-source Julia package and made publicly available on GitHub upon publication of this article.

## References

[1] Zhou, Y. et al. Membrane potential modulates plasma membrane phospholipid dynamics and k-ras signaling. Science 349, 873–876 (2015).

[2] Quang, B. A. T. et al. Extent of myosin penetration within the actin cortex regulates cell surface mechanics. Nature Communications 12, 6511 (2021).

[3] Svitkina, T. M. Actin cell cortex: Structure and molecular organization. Trends in cell biology 30, 556–565 (2020).

[4] Lelek, M. et al. Single-molecule localization microscopy. Nature Reviews Methods Primers 1, 39 (2021).

[5] Nieves, D. J. et al. A framework for evaluating the performance of SMLM cluster analysis algorithms. Nature Methods 20, 259–267 (2023).

[6] Pineda, J. et al. Enhanced spatial clustering of single-molecule localizations with graph neural networks. Nature Communications 16, 9693 (2025).

[7] Peters, R., Griffi, J., Burn, G. L., Williamson, D. J. & Owen, D. M. Quantitative fibre analysis of single-molecule localization microscopy data. Scientific Reports 8, 10418 (2018).

[8] Shirgill, S. et al. Nanoscale spatial-omics via contrastive embedding of single-molecule localisation data Preprint at bioRxiv 10.1101/2025.10.07.679170 (2025).

[9] Shirgill, S. et al. Nano-org, a functional resource for single-molecule localisation microscopy data. Nature Communications 16, 8674 (2025).

[10] Cooper, G. M. The cell: a molecular approach (ASM Press ; Sinauer Associates, 2000).

[11] Begg, D. A., Rodewald, R. & Rebhun, L. I. The visualization of actin filament polarity in thin sections. evidence for the uniform polarity of membrane-associated filaments. The Journal of Cell Biology 79, 846–52 (1978).

[12] Lschberger, A. et al. Super-resolution imaging visualizes the eightfold symmetry of gp210 proteins around the nuclear pore complex and resolves the central channel with nanometer resolution. Journal of Cell Science 125, 570–575 (2012).

[13] Jr., F. J. M. The Kolmogorov-Smirnov test for goodness of fit. Journal of the American Statistical Association 46, 68–78 (1951).

[14] Goodfellow, I. et al. Generative adversarial nets. Advances in Neural Information Processing Systems, Vol. 27, 2672–2680 (2014).

[15] Baniukiewicz, P., Lutton, J. E., Collier, S. & Bretschneider, T. Generative adversarial networks for augmenting training data of microscopic cell images. Frontiers in Computer Science 1, 10 (2019).

[16] Kingma, D. P. & Welling, M. Auto-encoding variational bayes. Preprint at arXiv 10.48550/arXiv.1312.6114 (2013).

[17] Ho, J., Jain, A. & Abbeel, P. Denoising diffusion probabilistic models. Advances in Neural Information Processing Systems, Vol. 33, 6840–6851 (2020).

[18] Saguy, A. et al. This microtubule does not exist: Super-resolution microscopy image generation by a diffusion model. Small Methods 9, 2400672 (2024).

[19] Bezanson, J., Edelman, A., Karpinski, S. & Shah, V. B. Julia: A fresh approach to numerical computing. SIAM Review 59, 65–98 (2017).

[20] Innes, M. et al. Fashionable modelling with Flux. Preprint at arXiv 10.48550/arXiv.1811.01457 (2018).

[21] Innes, M. Flux: Elegant machine learning with Julia. Journal of Open Source Software 3, 602 (2018).

[22] Mortenson, M. E. Mathematics for computer graphics applications (Industrial Press, 1999).

[23] Radosavovic, I., Kosaraju, R. P., Girshick, R., He, K. & Dollár, P. Designing Network Design Spaces. IEEE/CVF Conference on Computer Vision and Pattern Recognition (CVPR), 10428–10436 (2020).

[24] Loshchilov, I. & Hutter, F. Decoupled weight decay regularization. Preprint at arXiv 10.48550/arXiv.1711.05101 (2019).

[25] Smith, L. N. & Topin, N. Super-convergence: very fast training of neural networks using large learning rates. Proc. SPIE 11006, Artificial Intelligence and Machine Learning for Multi-Domain Operations Applications, 1100612 (2019).

