## Supplemental information for "SynthMLM: A framework for interpretable analysis and synthetic localisation data generation for SMLM"

### SynthMLM – Supplementary Information

This Supplementary Information provides additional use cases for SynthMLM together with a detailed description of the synthetic data generation framework, training data construction, image analysis model, the generation of descriptor-matched synthetic datasets, curvature reconstruction, and dose interpolation methodology.

#### 1 Clusters and Rings Analysis and Synthetic Data

SynthMLM is capable of dealing with clustered nanoscale localisation configurations (Figure S1A-C) and ring-like localisation patterns, e.g. nuclear pore complex [1] (Figure S1D-F).

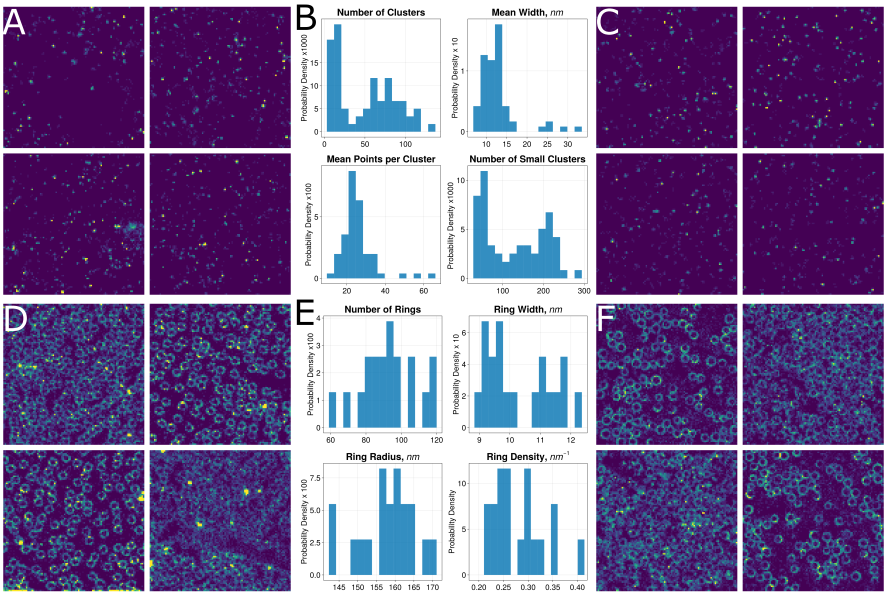


**Fig. S1** Example use case of SynthMLM with (top) clustered localisations and (bottom) ring-like structures (e.g. the nuclear pore complex, NPC). A) Whole-cell SMLM image of Lck in Jurkat E6.1 cells activated with anti-CD3/CD28 for 2 minutes. B) Normalised histograms of the inferred structural descriptors associated with clusters. 'Small clusters' refer to clusters containing $\leq$ 10 localisations, whereas 'clusters' refer to structures containing $>$ 10 localisations. C) Synthetic SMLM images generated using the inferred structural descriptors from the dataset shown in A. D) Whole-cell SMLM image of the Xenopus nuclear pore complex (NPC) [1], obtained from ShareLoc XYZ. E) Normalised histograms of the inferred structural descriptors associated with rings. F) Synthetic SMLM images generated using the inferred structural descriptors from the dataset shown in D.

### 2 Supplementary Methods

#### 2.1 Synthetic Data Generation

A synthetic SMLM image is built by generating zero or more structural elements (clusters, rings or fibres) consisting of randomly distributed particles (representing synthetic localisations) arranged around an underlying scaffold: points (clusters), circles (rings), or Bézier curves [2] (fibres). The placement of individual localisations is governed by multiple probability distributions, which are described below. These probability distributions correspond to real, measurable structural descriptors of an SMLM image.

Where possible, probability distributions are denoted by uppercase letters, and their realisations by lowercase letters. For example, if the number of particles in a cluster follows a distribution $D_C$, then the number of particles in the $i$th cluster is denoted by $d_{C,i}$. Unless otherwise stated, all parameters are expressed in physical units. Lengths are measured in nm, while curvature and densities are nm$^{-1}$.

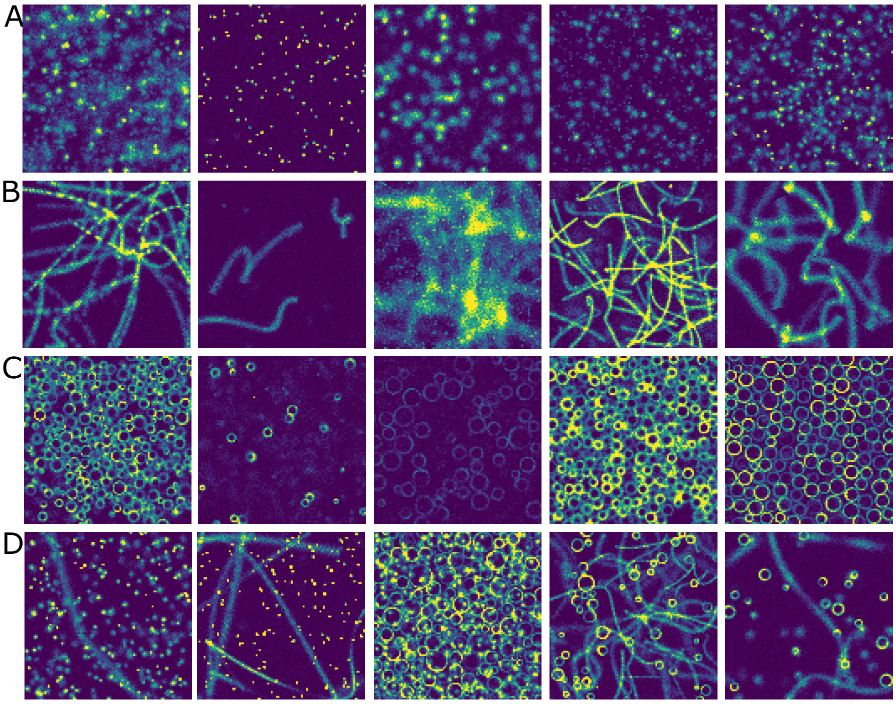


**Fig. S2** Representative synthetic SMLM images generated using SynthMLM showing A) clusters, B) fibres, C) rings, and D) mixtures of these structural elements (from left to right: clusters and fibres; small clusters and fibres; rings and clusters; fibres and rings; clusters, fibres, and rings.

Figure S2 illustrates the capabilities of the data generation aspect of SynthMLM, showing representative synthetic images of clusters, fibres, and ring-like structures. Although the full range of synthetic SMLM images generated by the software cannot be captured in a single figure, it can be seen that the synthetic data generation can cover a vast array of plausible (and implausible) biological conditions. Note that this is only a small sample of the possible images that SynthMLM can create, with over 30 parameters controlling the features of the generated elements (clusters, fibres and rings, with 4-10 parameters per element) and combinations thereof. Many generated images are intentionally biologically implausible, but these examples are retained during training to cover hypothetical and experimentally-unseen nanoscale configurations and widen the model's capabilities.

#### 2.2 Clusters

Currently, the software permits two different types of clusters in a given image, small clusters (with 10 or fewer particles per cluster), and large clusters (with more than 10 particles). These two types of clusters can have different structural descriptors to cover commonly seen experimental conditions: for example, to replicate localisation clusters with low occupancy clusters, or fibres breaking down into a mixture of small and large fragments.

The placement of clusters within an image is governed by three probability distributions for each cluster type: the number of clusters in an image, $N_Y$, the number of particles in a cluster $M_Y$ and the width of the clusters $\Sigma_Y$ for $Y = \{B, C\}$ for small and large clusters respectively. All clusters are Gaussian in shape, such that $\sigma$ is the standard deviation of its spatial distribution, with equal standard deviation in each dimension.

The $i$th cluster (with $i \in \{1, ..., n_C\}$) is defined by its centre, $c_i = (x_i, y_i)$, which is drawn from a uniform distribution in the image window, particle number $m_{C,i} \sim M_C$ and a width $\sigma_i \sim \Sigma_C$. The $j$th particle (for $j \in \{1, ..., m_{C,i}\}$) of cluster $i$ is then randomly placed at position $p_i^j \sim (x_i + N(0, \sigma_i), y_i + N(0, \sigma_i))$.

#### 2.3 Rings

A collection of $n_R$ rings has densities, $\lambda_{R,i}$, radii, $r_i$ and widths $w_{R,i}$, $i \in \{1, ..., n_R\}$, that are independently drawn for each ring from their respective distributions, $\Lambda_R$, $R$ and $W_R$. The radius of a ring is the distance from the centre to the middle of the annulus. The width refers to the spatial extent normal to the circle that defines the ring, i.e. the difference between the outer and inner radius of the annulus shape that defines the ring. The density $\lambda_{R,i}$ is measured along the ring's circumference, such that a ring with density $\lambda_{R,i}$ has $m_i = \text{round}(2\pi\lambda_{R,i}r_i)$ particles. At present, only a single ring type is allowed per image (i.e., rings are generated from a single tuple of distributions), reproducing the homogeneity in experimentally observed protein configurations like NPC [1].

A physics simulation is used to determine the spatial distribution of the rings. Initially, the ring centres are placed within the image window according to a uniform distribution. Any overlaps are resolved by modelling each ring as an inelastic circle and iteratively pushing overlapping rings apart. This process continues until the system relaxes and no overlaps remain. Only the radius is used in this physics simulation, ignoring the width of the rings.

In ring $i$, the position of particle $j$ (in polar coordinates, centred on the ring's centre) is selected as $r_i^j \sim r_i + N(0, \frac{1}{2}w_{R,i})$ where $r_i$ is the fixed radius and $w_{R,i}$ the width of ring $i$. The factor of $\frac{1}{2}$ is used to map the standard deviation of the normal distribution, centred along the middle of the ring, to what would be commonly understood as the width. The angular coordinate of particle $j$ is drawn from a von Mises distribution [3], to allow for angular inhomogeneity around the ring, recreating effects such as non-coplanarity with the imaging plane or uneven staining. This distribution is defined for each ring independently, with an angular location $\mu_i \sim U(0, 2\pi)$ and concentration $k_i \sim 2 \times \text{Beta}(2, 8)$. The distribution of $k_i$ was chosen heuristically to produce rings whose intensity profiles visually resemble experimentally observed data. When $k_i = 0$, the von Mises distribution reduces to the uniform distribution on $[0, 2\pi)$.

#### 2.4 Fibres

The fibres in an image are characterised by eight structural descriptors: $N_F$, which determines the number of fibres; $L$, the fibre length; $\Lambda_F$, the linear particle density; $W_F$, the fibre width; $\Delta$ and $\Xi$, which control the longitudinal inhomogeneity; and $B$ and $D$, which determine the geometric shape of the fibre.

An image is generated by first sampling the initial number of fibres $n_F^{\text{init}} \sim N_F$. This is the initial number of fibres the algorithm attempts to generate; due to random rotations and translations, fewer fibres may be captured in the image window. For each fibre $i \in \{1, \ldots, n_F^{\text{init}}\}$, independent realisations of the remaining parameters are drawn: $l_i$, $\lambda_{F,i}$, $w_{F,i}$, $\delta_i$, $\xi_i$, $b_i$ and $d_i$ in the same order as the distributions are listed in the previous paragraph.

##### 2.4.1 Shape

Each fibre is constructed by placing particles along a scaffold defined by a Bézier curve [2] of degree $d_i$. A Bézier curve of degree $d$ has $d + 1$ control points. For fibre $i$, the first and last control points are initially fixed at $(-l_i/2, 0)$ and $(l_i/2, 0)$ respectively. The remaining $d_i - 1$ control points are sampled independently from the region

$$S = \{(x, y) : x \in [-b_i l_i,\, b_i l_i],\ y \in [-b_i l_i,\, b_i l_i]\}, \tag{S1}$$

with $x \sim U(-b_i l_i, b_i l_i)$ and $y \sim U(-b_i l_i, b_i l_i)$.

These control points are then rotated around the origin by an angle $\phi$, and translated by a two dimensional vector $s_\mu$ (where the index $\mu$ is used for the two-dimensional

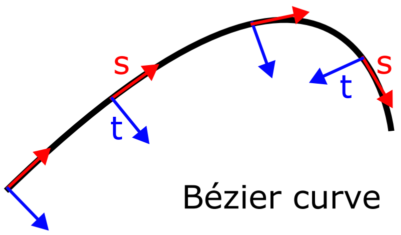


**Fig. S3** Parameterisation of particle positions around Bézier curve scaffold, showing the tangent coordinate $s$ in red and the normal coordinate $t$ in blue.

spatial components of vectors):

$$\phi \sim U(0, 2\pi) \qquad s_\mu \sim U(0, L_W), \quad \mu = 1, 2. \tag{S2}$$

Rotating and translating the control points is equivalent to rotating and translating the entire Bézier curve.

In the following section, we work with the arc-length parameterisation of these Bézier curves [4]. This parameter is denoted by $s \in [0, l_i]$ over the entire length of the fibre, although typically only a restricted subinterval, $s \in [s_0, s_1]$ (with $0 \leq s_0 < s_1 \leq l_i$) lies within the image window.

##### 2.4.2 Particle Distribution

To generate fibre $i$ with mean linear density $\lambda_i$, we place $m_i = \text{round}(\lambda_{F,i} l_i)$ particles around the Bézier curve scaffold. Particles are placed at randomly sampled fibre coordinates $(s_j, t_j)$, for $j \in \{1, ..., m_i\}$, where $s_j \in [0, l_i]$ is the tangential coordinate along the Bézier curve (in the arc-length parameterisation) and $t_j$ is the normal direction away from the curve, as shown in Figure S3. These coordinates are subsequently mapped to Cartesian coordinates in the image window when the particle's position is recorded.

To recreate the inhomogeneous appearance observed in real fibres, the particles are not uniformly distributed along the fibre, $s_j \nsim U(0, l_i)$. To achieve an inhomogeneous appearance, we build a unique probability distribution for the particle position along each fibre (the $s$ coordinate) in terms of a mixture of multiple normal distributions centred at randomly located intensity peaks. For each fibre, we sample $\delta_i \sim \Delta$, the distance between intensity peaks, and $\xi_i \sim \Xi$, the width of these intensity peaks. Let $m = \lfloor l_i/\delta_i \rfloor$ denote the approximate number of peaks along the fibre. To avoid boundary artefacts and to mimic fibres extending beyond the image window, we introduce $3m$ peaks indexed by $I \in \{-m, -m+1, \ldots, 2m\}$, so that an additional $m$ peaks lie in each of the extended regions $[-l_i, 0]$ and $[l_i, 2l_i]$.

To produce natural variation, we place the peaks along the fibre at

$$s_I^{\text{peak}} = I\delta_i + F_I + \delta_0, \tag{S3}$$

$$F_I \sim TN\left(0, \frac{\delta_i}{5}, -\frac{\delta_i}{2}, \frac{\delta_i}{2}\right),$$
$$\delta_0 \sim U\left(-\frac{\delta_i}{2}, \frac{\delta_i}{2}\right).$$

$F_I$ is a *per-peak* random shift to produce natural variation (truncated to preserve ordering and prevent peaks overlapping), and $\delta_0$ is a *per-fibre* offset to ensure that there is not always a peak at the beginning of the fibre.

The longitudinal coordinate $s_j$ of each particle is sampled from the normalised mixture with probability distribution function

$$S(s) = \frac{1}{Z} \sum_{I=-m}^{2m} N\left(s; s_I^{\text{peak}}, \xi_i\right), \qquad Z = \int_{-\infty}^{\infty} \sum_I N(s; s_I^{\text{peak}}, \xi_i)\, ds \tag{S4}$$

i.e. a normalised mixture of Gaussian distributions centred at the peak locations.

The transverse coordinate of particle $j$ is distributed according to the fibre width, $w_{F,i}$, $t_j \sim N(0, \frac{1}{2}w_{F,i})$.

To map fibre coordinates to Cartesian coordinates, the position of particle $j$ is defined by

$$\mathbf{p}_j = \mathbf{B}(s_j) + t_j \mathbf{N}_\mathbf{B}(s_j), \tag{S5}$$

where $\mathbf{B}(s) : \mathbb{R} \to \mathbb{R}^2$ is the Bézier curve parameterised by arc length and $\mathbf{N}_\mathbf{B}(s)$ is the corresponding unit normal vector at position $s$.

After this procedure, only the particles in the image window are retained, usually truncating the fibres. Additionally, if a fibre leaves the image window and returns at a later point, only the first portion that intersects the image window is kept. Output statistics, such as arc length and mean curvature, are computed using only the portion of each fibre contained within the image window.

#### 2.5 Final image

After all elements have been placed, the image is truncated to a square window of side length $L_W$, with vertices at $(0,0)$, $(L_W, 0)$, $(0, L_W)$, and $(L_W, L_W)$. In this work, we maintain the conventions established in nano-org, setting $L_W = 3\mu m$. The initial representation of an image is a two-dimensional array containing the coordinates of all particles within the window. For input into the image analysis model, this coordinate data is subsequently binned into a $100 \times 100$ pixel image, where each pixel corresponds to a square of side length 30 nm when $L_W = 3\mu m$.

#### 2.6 Training Data

We used the data generation software component of SynthMLM to generate 10 million images to be used as training data, featuring a curated mixture of different types of element, roughly proportional to the number of parameters needed to define each element. For example, there were far more images of fibres (alone and with clusters and rings, or both), because fibres involved multiple probability distributions and have a wide variety of shapes and configurations. Small clusters represent low-occupancy puncta, so they are present in all forms of images.

A training set of 10 million images was selected to balance diversity and computational practicality. The data was broken down as follows:

- 0.5 million × small clusters only,
- 1.5 million × clusters and small clusters,
- 2 million × fibres and small clusters,
- 2 million × clusters, fibres and small clusters,
- 1 million × rings and small clusters,
- 1 million × clusters, rings and small clusters,
- 1 million × fibres, rings and small clusters,
- 1 million × clusters, fibres, rings and small clusters.

For an image containing a given element type, the number of these elements is drawn from a discrete uniform distribution, with minimum and maximum values given in Table S1.

Each element has multiple structural descriptors that govern, e.g., their spatial extent, density and more. For the training data, these parameters are drawn from truncated normal distributions, $TN(\mu, s, l, u)$, defined by a mean $\mu$ and standard deviation $s$, and lower and upper bounds $l$ and $u$. $s$ is chosen to be a proportion of the mean value, $s = \mu r$. $\mu$ and $r$ are themselves drawn from continuous uniform distributions individually for each image. To summarise, for a parameter $X$ this is:

$$\begin{aligned} &X \sim TN(\mu_X, s_X, l_X, u_X), \qquad \mu_X \sim U(\alpha_X, \beta_X), \\ &s_X = \mu_X r_X, \quad r_X \sim U(q_X, p_X). \end{aligned} \tag{S6}$$

Here, $q_X$, $p_X$, $\alpha_X$, and $\beta_X$ are chosen for each parameter $X$ to span a broad range of plausible values, ensuring that the training data set captures as many biologically feasible conditions as possible. In the training data, $q_X = 0.1$ and $p_X = 0.5$ were used for all structural descriptors. $l_X$ and $u_X$ are selected to restrict possible structural descriptors to realistic and practically useful regimes. For example, these bounds ensure that physical quantities cannot be negative, that each individual element can be fully contained within the image window and remains visually distinguishable: rings must be visually distinguishable to avoid being misclassified as clusters, while fibres must be long enough to exhibit a clear fibrous morphology and not so wide or diffuse that this structure is lost.

There were two exceptions to this rule. First, $b$, which controls how far apart the control points of the Bézier curve can be placed, was fixed for all fibres in a given image. Second, $d$, the degree of a Bézier curve, was drawn from a discrete uniform distribution with a variable upper bound, chosen per image.

For every descriptor $X$ generated using this procedure, the corresponding values of $\alpha_X$, $\beta_X$, $l_X$, and $u_X$ are listed in Table S1. These values were determined through analysis of existing experimental data, supplemented by expert-guided extrapolation where necessary. Sometimes, $\alpha_X$, $\beta_X$, $l_X$ or $u_X$ are functions of related descriptors, e.g. $\delta$, the distance between peaks on the fibre is a function of the mean fibre length, $\mu_L$ (such that the length of fibres in a given image are $l \sim N(\mu_L, s_L)$).

For each image, the counts of each element and the mean and variance of the structural descriptor distributions visible in the image window are saved for all descriptors except the two parameters governing fibre shape, $b$ and $d$. Direct prediction of the Bézier parameters $b$ and $d$ offers little biological interpretability. Instead, these parameters are replaced by the mean curvature of the visible fibre segment, $\bar{\kappa}$, which is directly measurable from experimental data and has a clearer physical interpretation. When generating images, $b$ and $d$ are obtained from an empirical inverse function from the measured mean curvature, as outlined in Section 2.11.

For all elements, only the values of the structural descriptors visible in the image window are saved. For example, only the length of the fibres that are contained in the image window are recorded, along with all other fibre statistics, and the same for any rings that are pushed from the window due to the physics simulation. Fibres are only counted if the visible length is greater than twice their width, so that they look like a fibre. Rings are only counted if their centre lies within the image window. Due to the algorithm always placing clusters within the window, they are always counted.

We also built a test and validation set of 100 thousand images each, with the same relative proportions of image types as the training data.

#### 2.7 Preprocessing Data

For use with the model, we binned the particle positions into a $100 \times 100$ resolution image, with a single channel per pixel that represents the number of particles in that bin. These images were further pre-processed by dividing the pixel values by 40 (chosen as a generous upper-bound of the number of particles that could realistically occupy a $30\,\text{nm} \times 30\,\text{nm}$ region) and then clamping the values to lie between $[0, 1]$.

Regression labels were scaled using min-max normalisation. When reporting values to users, this scaling is reverted to restore the output variables to their physical units.

#### 2.8 Image Analysis Model

We built a convolutional neural network based on the RegNetX architecture [5]. RegNet models are composed of blocks grouped into stages, each stage systematically halves the resolution (using stride=2 convolutions) while increasing the channel width, $w$, by a factor of $\approx 2.5$ to compensate. A RegNet block is also defined by its group width, $g$ (chosen here such that there is always 16 groups) and bottleneck ratio, $k$, according to the best practice described in the original paper. The model design is outlined in Table S2.

#### 2.9 Training

To train the model, we used the AdamW optimiser [6] (with $\eta = 0.001, \alpha = 0.9, \beta = 0.99$ and $\lambda = 0.01$) and a OneCycle learning rate schedule [7], and a mean squared error loss function. We trained for 20 epochs and selected a final model using the validation set. The model was trained using an NVIDIA H100 GPU using the University of Birmingham's BlueBEAR HPC cluster.

To assess the predictive performance of the final image analysis model, we computed the coefficient of determination ($R^2$) for each output variable on the test set. The resulting scores are summarised in Table S3. The results demonstrate strong predictive performance across the majority of structural descriptors.

#### 2.10 Generating Descriptor-Matched Synthetic Datasets

To generate a descriptor-matched synthetic dataset from a given experimental dataset, the probability distributions of the descriptors that define the elements in the synthetic images are decided using two possible methods. After applying the image analysis model to each sub-image obtained from the dataset, we can either 1) treat these model outputs as sample means and variances of the descriptor distributions and sample from these distributions to generate synthetic data, or 2) generate synthetic data from inferred structural descriptors of randomly selected individual images from the original dataset. Method 1 was used for all synthetic data generation in the main paper, except for the synthetic datasets matching those from nano-org used in Figure 4, where Method 2 was used to capture the heterogeneity of the datasets.

##### 2.10.1 Method 1 – Dataset-level

The model is applied to every image in the original dataset to be replicated, $\mathcal{D}$, to produce a distribution for the counts of each element, and means and variances for every structural descriptor.

For the discrete counts, the distribution of possible numbers $N_Y$ of a given element $Y$ is a Poisson mixture model with each component Poisson distribution defined by a count from an image of $\mathcal{D}$. The final mixture distribution is truncated to lie between the minimum and maximum observed values of $N_Y$ in $\mathcal{D}$. For a dataset $\mathcal{D}$ with $n$

images, each containing (a model estimated) $\hat{N}_Y^i$ elements $Y$, this is (before rounding):

$$p(N_Y) = \frac{1}{n}\sum_{i=1}^{n} P(\hat{N}_Y^i) \tag{S7}$$

when $min(\hat{N}_Y^i) < N_Y < max(\hat{N}_Y^i)$ and 0 otherwise.

For the other structural descriptors, model outputs are treated as estimates of the sample means and variances of an underlying normal distribution. The true mean and variance of descriptor $X$ are estimated as the averages of the predicted sample means and variances across the dataset. Each descriptor $X$ is assumed to follow a truncated normal distribution with these means and variances, with $l_X$ and $u_X$ the same as the training data. Synthetic SMLM images are made by sampling these distributions and generating the resulting image as outlined above.

##### 2.10.2 Method 2 – Image-level

For some datasets, Method 1 is insufficient, particularly when the distribution of the structural descriptors does not follow a normal distribution: there may be two (or more) distinct spatial regions in a given dataset where the inferred descriptors differ, for example, fibres may be more numerous towards the middle of a cell. To better recreate this type of data, we can simply randomly select an image from $\mathcal{D}$ and generate a synthetic image based on the model output descriptor distributions from this single image. In doing so, we can generate data that better preserves the heterogeneity observed in the original dataset without interpolating between distinctly different regions.

#### 2.11 Curvature

When reconstructing synthetic data, each image is assigned a target mean curvature $\bar{\kappa}$ that it will attempt to reach. To do this, a probability distribution for $b$ and $d$ is calculated from the chosen value of $\bar{\kappa}$. To do this, we precomputed one million fibres spanning different values of $b$ and $d$, together with their corresponding mean curvatures. These samples form a lookup table used to estimate the conditional distribution of $b$ and $d$ for a given target mean curvature $\bar{\kappa}$, as can be seen in Figure S4.

Calling the collections of $b$, $d$ and mean curvatures $\bar{\kappa}$, $B_i$, $D_i$ and $\kappa_i$ respectively (with $i \in \{1, ..., 1000000\}$), we define:

$$I = \begin{cases} \{i : \bar{\kappa} - \epsilon \leq \kappa_i < \bar{\kappa} + \epsilon\} & \bar{\kappa} < 15 \\ \{i : \bar{\kappa} > 15\} & \bar{\kappa} \geq 15 \end{cases} \tag{S8}$$

where we used $\epsilon = 1/2$. The limit of 15 arises from considering Figure S4: after 15 the plot is thin for all values of $b$, reducing the set of possible values of $(b, d)$, so we accept any that created a mean curvature larger than 15. $I$ is the set of indices where

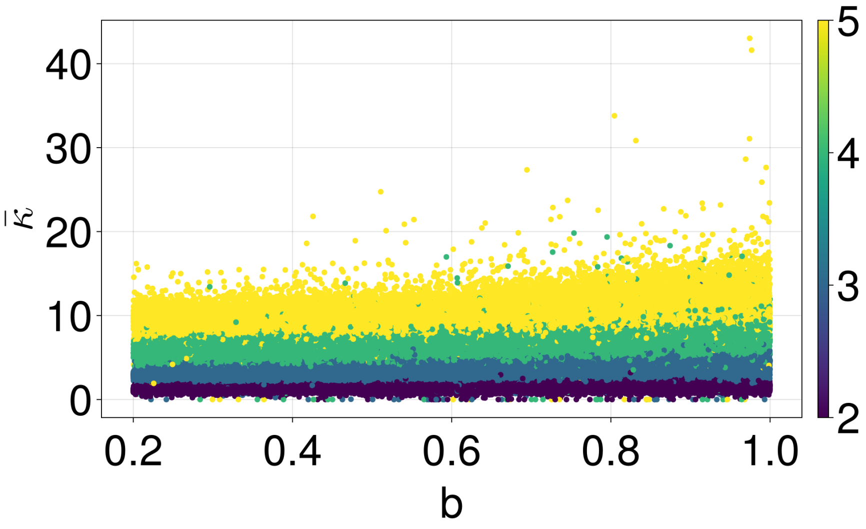


**Fig. S4** Mean curvature, $\bar{\kappa}$ against $b$ (x-axis) and $d$ (colour) for 1,000,000 generated curves used to define inverse function $b(\bar{\kappa})$ and $d(\bar{\kappa})$.

the curvature falls in a pre-defined range, to obtain a $b$ and a $d$ we randomly sample from $B_i$ and $D_i$ with $i \in I$.

#### 2.12 Interpolation of Experimental Conditions

For Figure 5 in the main manuscript, we used 3 existing doses to interpolate the effects of a fourth dose. This was done by first taking the mean of every model output descriptor across each condition, providing a triple of values for each model output, $\bar{X}^i_j$ for $i \in \{1, ..., 30\}$ and $j \in \{0, 0.1, 1\}$ $\mu$g/mL. Consistent with conventional models of drug dose-response relationships, we assumed each $\bar{X}^i$ lay on a Hill curve where

$$\bar{X}^i(D) = \begin{cases} A^i \frac{K^i}{D^{n^i}+K^i} + B^i & \bar{X}^i(0) > \bar{X}^i_1 \\ A^i \frac{D^{n^i}}{D^{n^i}+K^i} + B^i & \bar{X}^i(0) \leq \bar{X}^i_1 \end{cases} \tag{S9}$$

For each structural descriptor $X$, the values of $A^i$, $B^i$ and $K^i$ were fit by the requirements that $\bar{X}^i(0) = \bar{X}^i_0$ and $\bar{X}^i(1) = \bar{X}^i_1$, and to minimise the distance between the predicted $\bar{X}^i(0.1)$ and the measured $\bar{X}^i_{0.1}$ using a least squares estimate. We chose $n^i = 1$ for simplicity, but this could also be estimated from the data. In doing so we built a simple function of dose dependency on each input descriptor to the synthetic data generator, which was used to determine $\bar{X}^i(0.5\,\mu\text{g/mL})$ and generate a hypothetical synthetic representation of the $0.5\,\mu$g/mL nocodazole condition.

### 3 Supplementary Tables

**Table S1** Parameter distributions governing the generation of images in the training data for the SMLM image analysis model.

| Descriptor | Description | Units | $\mu$ | $l$ | $u$ |
|---|---|---|---|---|---|
| Clusters | | | | | |
| $N_C \sim \mathcal{U}(l,u)$ | Number of clusters | N/A | N/A[1] | 1 | 800 |
| $m_C \sim M_C$ | Particles per cluster | N/A | $U(11, 400)$ | 11 | 1000 |
| $\sigma_C \sim \Sigma_C$ | Width of clusters | nm | $U(5, 100)$ | 5 | 300 |
| Small clusters | | | | | |
| $N_B \sim \mathcal{U}(l,u)$ | Number of small clusters | N/A | N/A[1] | 1 | 500 |
| $m_B \sim M_B$ | Particles per small cluster | N/A | $U(1, 10)$ | 1 | 50 |
| $\sigma_B \sim \Sigma_B$ | Width of small clusters | nm | $U(5, 60)$ | 5 | 180 |
| Rings | | | | | |
| $N_R \sim \mathcal{U}(l,u)$ | Number of rings | N/A | N/A[1] | 1 | 400 |
| $r \sim R$ | Radius of rings | nm | $U(55, 120)$ | 25 | 300 |
| $w_R \sim W_R$ | Width of rings | nm | $U(10, W_0)$[2] | 6 | $3W_0$ |
| $\lambda_R \sim \Lambda_R$ | Density of rings | $nm^{-1}$ | $U(0.2, 1.5)$ | 0.1 | 4.5 |
| Fibres | | | | | |
| $N_F \sim \mathcal{U}(l,u)$ | Number of fibres | N/A | N/A[1] | 1 | 50 |
| $l \sim L$ | Length of fibres | nm | $U(1000, 6000)$ | 500 | 10000 |
| $\lambda_F \sim \Lambda_F$ | Density of fibres | $nm^{-1}$ | $U(0.2, 2)$ | 0.25 | 6 |
| $w_F \sim W_F$ | Width of fibres | nm | $U(20, 200)$ | 10 | 600 |
| $\delta \sim \Delta$ | Distance between fibre intensity peaks | nm | $U(\mu_L/40, \mu_L/4)$ | $\mu_L/100$ | $\mu_L/2$ |
| $\xi \sim \Xi$ | Width of fibre intensity peaks | nm | $U(\mu_\Delta/4, \mu_\Delta)$ | $\mu_\Xi/2$ | $3\mu_\Delta$ |
| $d \sim \mathcal{U}(l,u)$ | Maximum degree of Bézier curve | N/A | N/A | 2 | $\mathcal{U}(2, 5)$ |
| $b \sim U(l,u)$ [3] | Bézier curve control point range | N/A | N/A | 0.2 | 1 |

[1] The number of a particular element is discrete uniformly distributed in the training data.

[2] $W_0 = min(40, 14r/30)$, determined heuristically: to ensure rings have a visible hole, the width had to be above $14r/30$ for a ring of radius $r$.

[3] In the training data, $b$ was fixed for all fibres in an image.

**Table S2** Network structure of the image analysis model aspect of SynthMLM. When the width and resolution doesn't change in a block (or chain of blocks), a single value is given, but if the width or resolution is changed, we write $(x, x) \rightarrow (y, y)$ for input width/resolution $(x, x)$ and output width/resolution $(y, y)$. $\times X$ means the same convolution is performed $X$ times sequentially. Block refers to a RegNet block, which is a stack of three convolutions and a residual connection. After each convolution there is a BatchNorm followed by ReLu.

| Layer | Resolution | $w$ | $g$ | $k$ |
|---|---|---|---|---|
| Stem | | | | |
| $\text{Conv}_{s=2}$ | $(100, 100) \rightarrow (50, 50)$ | $1 \rightarrow 46$ | 1 | 1 |
| Body: Stage 1 | | | | |
| $\text{Block}_{s=2}$ | $(50, 50) \rightarrow (25, 25)$ | $46 \rightarrow 128$ | 8 | 1 |
| Block $\times 3$ | $(25, 25)$ | 128 | 8 | 1 |
| Body: Stage 2 | | | | |
| $\text{Block}_{s=2}$ | $(25, 25) \rightarrow (13, 13)$ | $128 \rightarrow 320$ | 20 | 1 |
| Block $\times 4$ | $(13, 13)$ | 320 | 20 | 1 |
| Body: Stage 3 | | | | |
| $\text{Block}_{s=2}$ | $(13, 13) \rightarrow (8, 8)$ | $320 \rightarrow 800$ | 50 | 1 |
| Block $\times 5$ | $(8, 8)$ | 800 | 50 | 1 |
| Body: Stage 4 | | | | |
| $\text{Block}_{s=2}$ | $(8, 8) \rightarrow (4, 4)$ | $800 \rightarrow 2000$ | 125 | 1 |
| Block $\times 3$ | $(4, 4)$ | 2000 | 125 | 1 |
| Head | | | | |
| AdaptiveMeanPool(1,1) | $(4, 4) \rightarrow (1, 1)$ | 2000 | N/A | N/A |
| Dense(2000, 30) | N/A | N/A | N/A | N/A |

**Table S3** Coefficient of determination ($R^2$) between the inferred and true values of each predicted output for the final selected model on the test dataset. Results are reported for element counts and the mean and variance of each structural descriptor. See Table S1 for the definition of each descriptor.

| Descriptor | Count/Mean $R^2$ | Variance $R^2$ |
|---|---|---|
| Clusters | | |
| $N_C$ | 0.974 | N/A |
| $m_C$ | 0.962 | 0.865 |
| $\sigma_C$ | 0.949 | 0.923 |
| Small clusters | | |
| $N_B$ | 0.630 | N/A |
| $m_B$ | 0.584 | 0.428 |
| $\sigma_B$ | 0.567 | 0.396 |
| Rings | | |
| $N_R$ | 0.995 | N/A |
| $r$ | 0.991 | 0.978 |
| $w_R$ | 0.985 | 0.946 |
| $\lambda_R$ | 0.994 | 0.969 |
| Fibres | | |
| $N_F$ | 0.957 | N/A |
| $l$ | 0.940 | 0.856 |
| $w_F$ | 0.948 | 0.898 |
| $\lambda_F$ | 0.959 | 0.843 |
| $\delta$ | 0.733 | 0.585 |
| $\xi$ | 0.705 | 0.572 |
| $\bar{\kappa}$ | 0.653 | 0.633 |

### References


[1] Lschberger, A. *et al.* Super-resolution imaging visualizes the eightfold symmetry of gp210 proteins around the nuclear pore complex and resolves the central channel with nanometer resolution. *Journal of Cell Science* **125**, 570–575 (2012).

[2] Mortenson, M. E. *Mathematics for computer graphics applications* (Industrial Press, 1999).

[3] Mardia, K. V. & Jupp, P. E. *Directional statistics* (J. Wiley, 2000).

[4] do Carmo, M. *Differential Geometry of Curves and Surfaces: Revised and Updated Second Edition* Dover Books on Mathematics (Dover Publications, 2016).

[5] Radosavovic, I., Kosaraju, R. P., Girshick, R., He, K. & Dollár, P. Designing Network Design Spaces. *IEEE/CVF Conference on Computer Vision and Pattern Recognition (CVPR)*, 10428–10436 (2020).

[6] Loshchilov, I. & Hutter, F. Decoupled weight decay regularization. Preprint at *arXiv* https://doi.org/10.48550/arXiv.1711.05101 (2019).

[7] Smith, L. N. & Topin, N. Super-convergence: very fast training of neural networks using large learning rates. *Proc. SPIE 11006, Artificial Intelligence and Machine Learning for Multi-Domain Operations Applications, 1100612* (2019).
